# A two-component signature determines which rust fungi secreted proteins are translocated into the cells of the host plant

**DOI:** 10.1101/2024.07.01.601607

**Authors:** Gregory J Lawrence

## Abstract

Rust diseases of plants are caused by parasitic fungi that feed off living plant cells by means of haustoria that form within plant cells. These haustoria also secrete a large number of proteins, some of which remain in the matrix surrounding the haustoria while others are translocated through a membrane into the cytoplasm of the plant cell. These latter proteins would be expected to possess a signature marking them out for translocation but, to date, no such signature has been identified. An examination of a set of wheat rust proteins known to be translocated to the cytoplasm of the wheat cell, together with an analysis of 1208 wheat stem rust (*Puccinia graminis* f. sp. *tritici*) secretome proteins, provides evidence that the translocation signature contains two components. The first component consists of a positively-charged amino acid at position 1, 2 or 3 (and possibly 4 or greater) upstream of the hydrophobic region in the signal peptide. The second component consists of a positively-charged amino acid at position 21 downstream of the signal peptide. A similar analysis of flax rust (*Melampsora lini*) secretome proteins indicates that the first component is the same as that of the wheat stem rust secretome proteins but that the second component consists of a positively-charged amino acid at position (16)17-20 downstream of the signal peptide. The flax rust signature may also be employed by wheat stem rust in its pycnial stage when growing on its alternate dicot host, barberry. Being able to identify which rust haustorial secreted proteins go to the plant cytoplasm and which to the extrahaustorial matrix should facilitate work aimed at identifying the function of each of the secreted proteins and, also, work aimed at elucidating the translocation mechanism, an understanding of which could open up new approaches to rust control.

## Introduction

Rust fungi are biotrophic plant pathogens that obtain nutrients from living plant cells via haustoria that form within plant cells after the fungus gains entry to the cell through a small hole that it makes in the cell wall. These haustoria, which have their own cell wall and internal membrane, are separated from the cytoplasm of the plant cell by an extrahaustorial matrix layer and an extrahaustorial membrane which, although an extension of the plasma membrane of the plant cell, is newly formed and therefore could have a different lipid and protein composition to that of the plasma membrane, potentially under the control of the rust pathogen.

While nutrient uptake is a major role of rust haustoria, a second role, of equal importance, is that of protein secretion [1,2]. Proteins secreted by haustoria either enter the extrahaustorial matrix or are translocated to the cytoplasm of the plant cell by an unknown mechanism. It would be expected that these latter proteins would carry a signature of some kind marking them out for translocation. However, no conserved motif has been identified in rust proteins known to be translocated into the plant cell, unlike in the oomycete pathogens, such as the downy mildews and *Phytophthora* species, where an Arg-Xaa-Leu-Arg (RxLR) motif near the N-terminus of a protein marks that protein for translocation [3,4].

This paper reports observations that indicate that the translocation signature for rust secreted proteins contains two components – one located in the signal peptide of the protein and the other in the N-terminus region of the protein.

## Methods

The amino acid sequences of secretome proteins from wheat stem rust, *Puccinia graminis* f. sp. *tritici*, and flax rust, *Melampsora lini*, were submitted to the signal peptide predictors PHOBIUS [5] and SIGNALP 6.0 [6] to identify for each protein the length of its signal peptide and the N-terminal, hydrophobic and C-terminal subregions of the signal peptide. The position upstream of the hydrophobic region of any positively-charged amino acids in the N-terminal region was noted as were the positions of any positively-charged amino acids in the first 30 amino acids downstream of the signal peptide. The relationship between the presence of a positively-charged amino acid at particular positions in the N-terminal region of the signal peptide and the frequency of positively-charged amino acids at positions downstream of the signal peptide was examined. The signal peptide predictors SIGNALP 4.1 [7,8] and SIGNALP 5.0 [9] were also employed on specific occasions.

## Results

In the search for a translocation signature in rust secreted proteins, 11 rust secreted proteins known to enter the plant cell were examined for any feature(s) that they might have in common. Of these 11, seven (AvrL567-A, AvrL2-A, AvrM-A, AvrM14-A, AvrP, AvrP123-A and AvrP4) are avirulence proteins from flax rust [10,11,12,13], two (AvrSr35 and AvrSr50) are avirulence proteins from wheat stem rust [14,15], while the remaining two, PGTG-08638 = PGTAUSPE10 from wheat stem rust and Pst_8713 from wheat stripe rust, *Puccinia striiformis* f. sp. *tritici*, have both been shown to enter the plant cell [16,17,18]. The avirulence proteins are assumed to enter the plant cell because they are recognised by plant resistance gene proteins that are located inside plant cells. Direct protein interaction between AvrSr35, AvrSr50, AvrL567-A and AvrM-A and their corresponding host resistance proteins (immune receptors) has been demonstrated [14,15,19,20].

The signal peptides of the tester set of proteins were examined using the PHOBIUS signal peptide predictor [5]. This predictor, where it predicts the presence of a signal peptide, also predicts the N-terminal, hydrophobic and C-terminal regions of each signal peptide. PHOBIUS did not identify a signal peptide in one, AvrP123-A, of the proteins in the tester set but, as shown in Fig. 1, a feature that all 10 of the other signal peptides have in common is that each possesses a positively-charged amino acid, either arginine (R), lysine (K) or histidine (H), at position 2 or position 3 upstream of the hydrophobic region in the N-terminal region. An examination of 524 flax rust secretome proteins (see below) found that 28.05% of them possess a positively-charged amino acid at position 2 and/or 3 in the N-terminal region. Therefore the probability that six randomly-chosen flax rust secretome proteins would all possess this feature is (0.2805)^6^ or 0.000487. Likewise, an examination of the signal peptides of 1208 wheat stem rust secreted proteins (see below) found that 29.9% had a positively-charged amino acid at position 2 and/or 3 in their N-terminal region. Therefore the probability that four randomly-chosen wheat rust secretome proteins would all possess this feature is (0.299)^4^ or 0.0079925 (assuming the single wheat stripe rust protein has the same probability). Combined, the probability that all 10 proteins would possess a positively-charged amino acid at positions 2 and/or 3 just by chance is the product of the two separate probabilities which is 0.0000038: this is such a small probability that the possibility that chance gave rise to this common feature in the tester set of proteins can be rejected. Therefore, this common feature, the presence of a positively-charged amino acid at position 2 and/or 3 in the N-terminal region of a signal peptide, can be considered a putative translocation signature.

**Figure 1.**
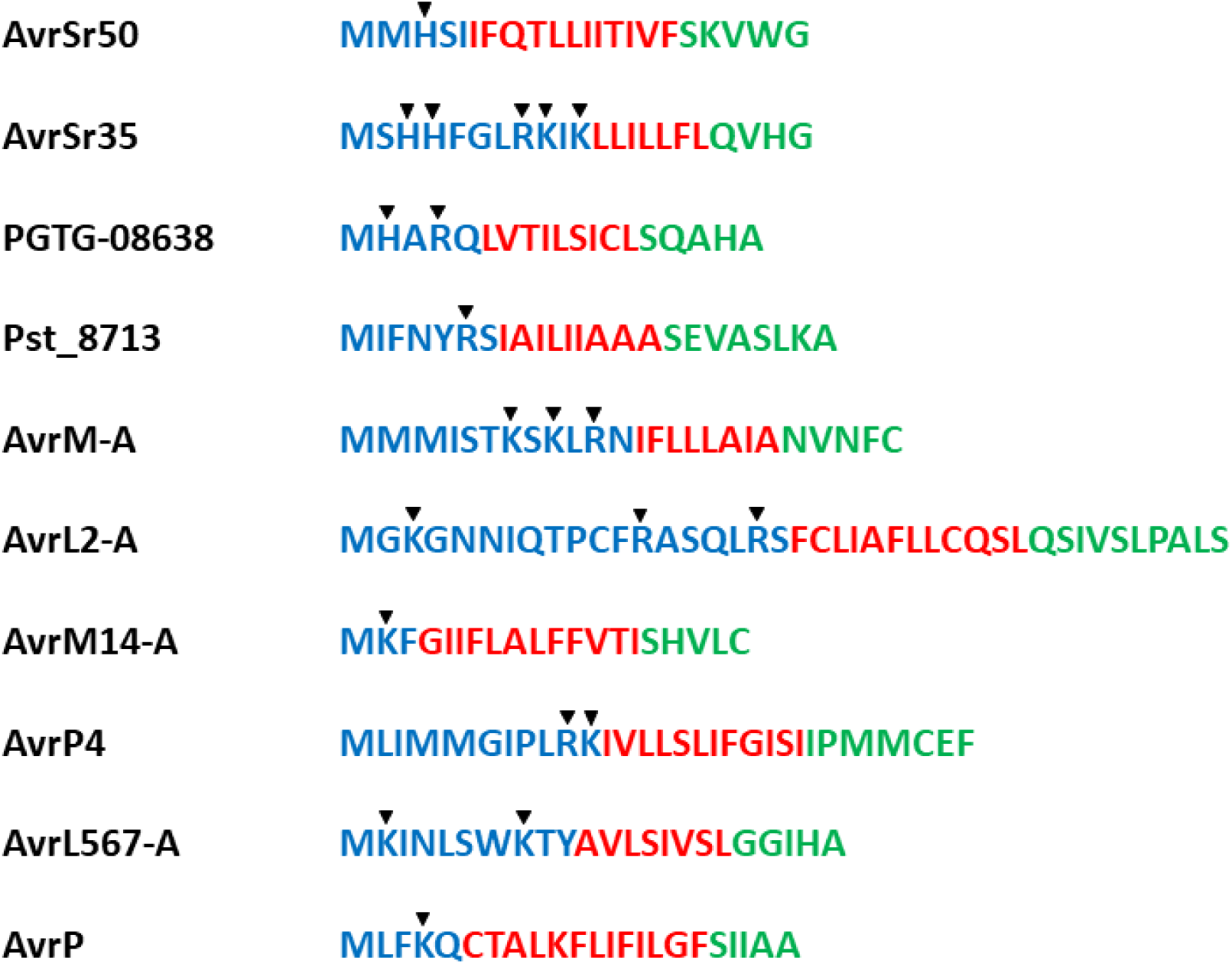
The amino acid sequences of the signal peptides, as predicted by PHOBIUS, of wheat stem rust, wheat stripe rust and flax rust secretome proteins that are known to be translocated to the cytoplasm of the plant cell. The N-terminal region is shown in blue, the hydrophobic region in red and the C-terminal region in green. Arrowheads indicate positively-charged amino acids.

Of the 1208 wheat stem rust secretome proteins examined, 361 possess a positively-charged amino acid at positions 2 and/or 3 in the N-terminal region of their signal peptides (see below). As it seems unlikely that this number of proteins would be translocated into the plant cell, a second feature common to all 11 proteins in the tester set was searched for. An examination of the location of positively-charged amino acids downstream of the signal peptide revealed that all four wheat rust proteins possess a positively-charged amino acid at position 21, while the flax rust proteins all possess a positively-charged amino acid in the 17 to 20 region (Fig. 2). Largely because the four wheat rust proteins are identical with respect to this character, this character was tentatively noted as a putative signature. The probability that the four wheat rust proteins would be identical for this character by chance alone is (0.1118)^4^ or 0.000156 since, amongst 1208 wheat stem rust secretome proteins 135 (11.18%) were found to possess a positively-charged amino acid at position 21 downstream of the signal peptide (see below). The probability calculation above assumes that the single wheat stripe rust protein has the same probability (0.1118) as the wheat stem rust proteins.

**Figure 2.**
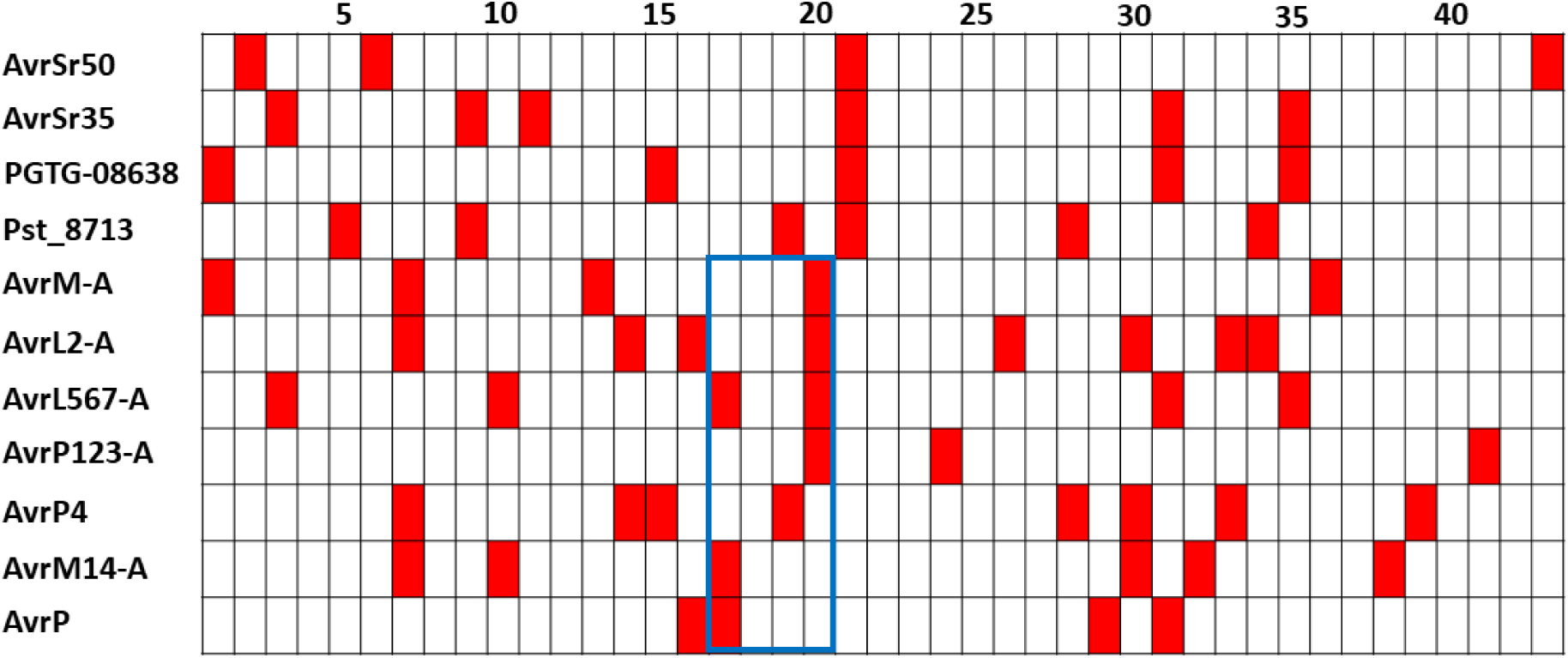
The location of positively-charged amino acids (filled boxes) downstream of the signal peptide (as determined by SIGNALP 4.1) of 11 wheat and flax rust secretome proteins that are known to be translocated to the cytoplasm of the plant cell. The blue rectangle frames amino acids at positions 17 to 20 downstream of the signal peptides of the flax rust proteins.

In Fig. 2 the signal peptide size predicted by another predictor, SIGNALP 4.1 [7,8], has been used which results in all four wheat rust proteins having a positively-charged amino acid at position 21 downstream of the signal peptide. The PHOBIUS prediction for the length of the signal peptide of AvrSr35 (22 amino acids) differs from that of SIGNALP 4.1 (25 amino acids) – consequently only three of the four wheat rust proteins would have a positively-charged amino acid at position 21 if the PHOBIUS size predictions had been used.

The availability of a list of wheat stem rust secretome proteins that do not possess a transmembrane domain [21] provided an opportunity to investigate if there might be a relationship between the putative signal peptide signature and the presence of a positively-charged amino acid at position 21. If both of these putative signatures are required for translocation then, in a group of proteins with the signal peptide N-terminal signature, the frequency with which a positively-charged amino acid occurs at position 21 will depend on how many in the group are targeted for translocation to the cytoplasm of the plant cell. However, the frequency with which a positively-charged amino acid occurs at other positions will depend on how often a positively-charged amino acid at a particular position contributes to the functioning of the protein of which it is part. These frequencies could differ. Therefore, in a group of proteins with the putative signal peptide signature, if the frequency of a positively-charged amino acid at position 21 differs markedly from that at the surrounding positions, this would support the case for a positively-charged amino acid at position 21 being a translocation signature when the putative signal peptide signature is also present.

For this analysis a list of 1924 wheat stem rust secretome proteins identified using SIGNALP 4.0 following the sequencing of the wheat stem rust genome [21, Supplementary Table S7] was used. The N-terminal region of the signal peptide as predicted by PHOBIUS [5] and the length of the signal peptide as predicted by SIGNALP 6.0 (long output, slow model mode) [6] were used in the analysis. SIGNALP 6.0 was used because, unlike previous versions of SIGNALP, it also identifies the N-terminal, hydrophobic and C-terminal regions of the signal peptide and it was of interest to compare its predictions for the sub-regions to those of PHOBIUS.

From the original list of 1924 secretome proteins that were identified using SIGNALP 4.0, 314 were omitted because they did not have a PASA (Program to Assemble Spliced Alignments) pass. Of the remainder (1610), 133 were excluded because their amino acid sequence for the first 70 amino acids was observed to be identical to that of another protein adjacent to, or nearby, in the list. Also excluded were another 269 proteins for which PHOBIUS and/or SIGNALP 6.0 did not predict the presence of a signal peptide. This left 1208 proteins for the analysis.

Table 1 shows the frequency of positively-charged amino acids at the first 30 positions downstream of the signal peptide of 1208 wheat stem rust secretome proteins that have been divided into seven groups based on the positively-charged amino acid composition of the N-terminal region of their signal peptide. Amongst 379 proteins that possess no positively-charged amino acids in the signal peptide N-terminal region (Table 1, row 1), 52 possess a positively-charged amino acid at position 21. This score sits well within the range of scores (39 to 68) observed for positions 5 to 30, with 13 positions having a lower score and 12 a higher score. Fig. 3A shows the results in bar chart form.

**Figure 3.**
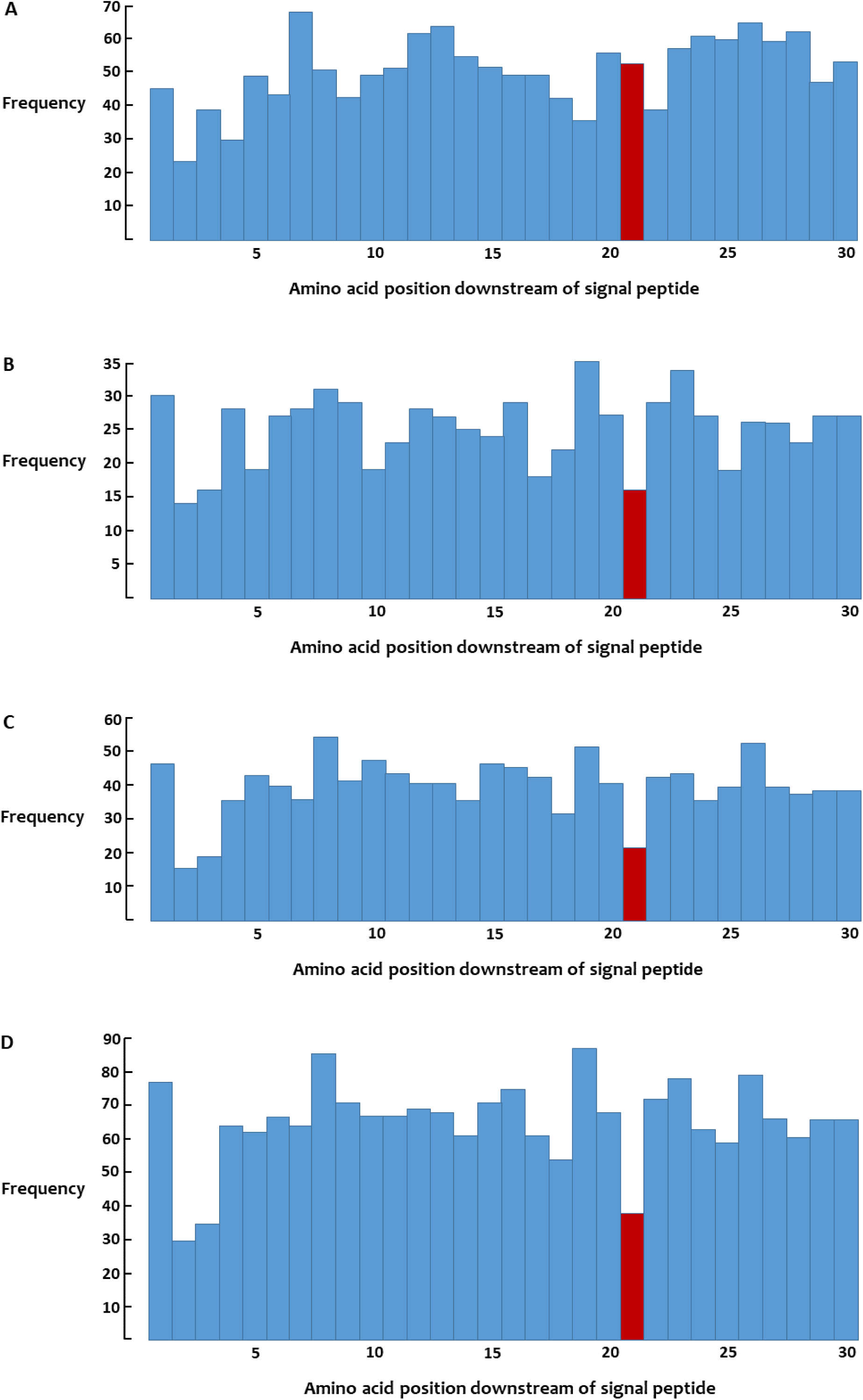
Frequency of positively-charged amino acids at positions 1 to 30 downstream of the signal peptide of wheat stem rust secretome proteins. **A**. Group of 379 proteins that have no positively-charged amino acids upstream of the hydrophobic region in their signal peptides. **B**. Group of 194 proteins that have a single positively-charged amino acid at either position 2 or position 3 upstream of the hydrophobic region in their signal peptides. **C**. Group of 293 proteins that have a single positively-charged amino acid at position 1 upstream of the hydrophobic region in their signal peptides. **D**. Group of 487 proteins that have a single positively-charged amino acid at either position 1 or position 2 or position 3 upstream of the hydrophobic region in their signal peptides.

**Table 1.** Frequency of positively-charged amino acids at positions 1 to 30 downstream of the signal peptide of 1208 wheat stem rust secretome proteins grouped according to their positively-charged amino acid composition in the N-terminal region of their signal peptides

| Row | 1 | 2 | 3 | 4 | 5 | 6 | 7 | 8 | 9 | 10 | 11 | 12 | 13 | 14 | 15 | 16 | 17 | 18 | 19 | 20 | 21 | 22 | 23 | 24 | 25 | 26 | 27 | 28 | 29 | 30 |
| --- | --- | --- | --- | --- | --- | --- | --- | --- | --- | --- | --- | --- | --- | --- | --- | --- | --- | --- | --- | --- | --- | --- | --- | --- | --- | --- | --- | --- | --- | --- |
| 1 | 45 | 24 | 39 | 30 | 49 | 43 | 68 | 51 | 43 | 49 | 51 | 62 | 64 | 55 | 51 | 49 | 49 | 42 | 36 | 56 | <u>52</u> | 39 | 57 | 61 | 60 | 65 | 59 | 62 | 47 | 53 |
| 2 | 47 | 16 | 19 | 36 | 43 | 40 | 36 | 55 | 42 | 48 | 44 | 41 | 41 | 36 | 47 | 46 | 43 | 32 | 52 | 41 | <u>22</u> | 43 | 44 | 36 | 40 | 53 | 40 | 38 | 39 | 39 |
| 3 | 15 | 7 | 10 | 16 | 11 | 18 | 16 | 20 | 17 | 16 | 14 | 19 | 19 | 19 | 17 | 18 | 10 | 13 | 23 | 19 | <u>11</u> | 18 | 21 | 22 | 15 | 18 | 16 | 16 | 22 | 18 |
| 4 | 15 | 7 | 6 | 12 | 8 | 9 | 12 | 11 | 12 | 3 | 9 | 9 | 8 | 6 | 7 | 11 | 8 | 9 | 12 | 8 | <u>5</u> | 11 | 13 | 5 | 4 | 8 | 10 | 7 | 5 | 9 |
| 5 | 77 | 30 | 35 | 64 | 62 | 67 | 64 | 86 | 71 | 67 | 67 | 69 | 68 | 61 | 71 | 75 | 61 | 54 | 87 | 68 | <u>38</u> | 72 | 78 | 63 | 59 | 79 | 66 | 61 | 66 | 66 |
| 6 | 6 | 6 | 13 | 3 | 14 | 14 | 15 | 12 | 10 | 18 | 17 | 12 | 14 | 16 | 15 | 9 | 8 | 9 | 16 | 12 | <u>18</u> | 12 | 9 | 13 | 10 | 12 | 16 | 10 | 21 | 13 |
| 7 | 14 | 14 | 8 | 16 | 22 | 12 | 15 | 12 | 19 | 13 | 12 | 16 | 15 | 9 | 16 | 20 | 24 | 24 | 18 | 14 | <u>13</u> | 20 | 20 | 10 | 21 | 15 | 16 | 16 | 18 | 12 |
| 8 | 97 | 50 | 56 | 83 | 98 | 93 | 94 | 110 | 100 | 98 | 96 | 97 | 97 | 86 | 102 | 104 | 93 | 87 | 121 | 94 | <u>69</u> | 104 | 107 | 86 | 90 | 106 | 98 | 87 | 105 | 91 |
| 9 | 14 | 12 | 10 | 13 | 16 | 19 | 13 | 18 | 14 | 16 | 12 | 22 | 10 | 14 | 15 | 16 | 18 | 15 | 14 | 17 | <u>14</u> | 22 | 13 | 11 | 14 | 14 | 13 | 8 | 16 | 15 |

Row 1: No positively-charged amino acids in the N-terminal region of the signal peptide – group of 379 proteins
Row 2: Single positively-charged amino acid at position 1 upstream of hydrophobic region of signal peptide – group of 293 proteins
Row 3: Single positively-charged amino acid at position 2 upstream of hydrophobic region of signal peptide – group of 135 proteins
Row 4: Single positively-charged amino acid at position 3 upstream of hydrophobic region of signal peptide – group of 59 proteins
Row 5: Sum of rows 2, 3 and 4 – group of 487 proteins
Row 6: Two or three positively-charged amino acids at positions 1, 2 and 3 upstream of hydrophobic region of signal peptide with or without additional positively-charged amino acids at position 4 or greater – group of 107 proteins
Row 7: Single positively-charged amino acid at either position 1, 2 or 3 upstream of hydrophobic region of signal peptide together with one or more positively-charged amino acids at position 4 or greater – group of 129 proteins
Row 8: Sum of rows 5, 6 and 7 – group of 723 proteins
Row 9: No positively-charged amino acid at position 1, 2 or 3 upstream of hydrophobic region of signal peptide but have one or more positively-charged amino acids at position 4 or greater – group of 106 proteins

Considering those proteins with a putative signature, amongst 135 proteins with a single positively-charged amino acid at position 2 in the N-terminal region of their signal peptides (Table 1, row 3), 11 possess a positively-charged amino acid at position 21 which is right at the lower end of the range of scores observed (10 to 22) for positions 5 to 30, with only one position having a lower score. Amongst 59 proteins with a single positively-charged amino acid at position 3 in the N-terminal region of the signal peptide the score for position 21 is 5, which is again at the lower end of the range (3 to 12) for positions 5 to 30 with only two positions having a lower score than position 21 (Table 1, row 4). When these two groups with putative signature signal peptides are combined, the score for position 21 is 16, which is lower than the combined score for any of the other (5 to 30) positions (range 18 to 35, Fig. 3B).

The number of proteins with a single positively-charged amino acid at position 1 in the N-terminal region of the signal peptide (293) is much larger than the number in the position 2 and 3 groups (135 and 59 respectively) considered above. With this larger group the score for position 21 is 22, which is notably lower than the range of 32 to 55 amongst positions 5 to 30 (Table 1, row 2; Fig. 3C). So position 21 clearly stands out in this group, suggesting that a positively-charged amino acid at position 1 in the N-terminal region of the signal peptide is also a signature component. When proteins containing a single positively-charged amino acid at either positions 1, 2 or 3 in the N-terminal region of the signal peptide are combined, the score for position 21 is 38, which is markedly lower than that for any of the other (5 to 30) positions which range from 54 to 87 (Table 1, row 5; Fig. 3D).

Some proteins contained two or three positively-charged amino acids at positions 1, 2 or 3 in the N-terminal region of their signal peptides with or without additional positively-charged amino acids at positions 4 or greater in the N-terminal region of the signal peptide. Interestingly, in this group of 107 proteins, the positively-charged amino acid score for position 21 is 18, which places it in the second highest position in the range of 8 to 21 amongst positions 5 to 30 (Table 1, row 6). Thus having two or three positively-charged amino acids in the putative signature positions in the N-terminal region is apparently associated with a relatively higher score at position 21. One of the wheat stem rust proteins, AvrSr35, (Fig. 1) is in this category.

There are also proteins that have a positively-charged amino acid at either position 1, 2 or 3 in the N-terminal region of the signal peptide together with one or more positively-charged amino acids at position 4 or greater (Table 1, row 7). When combined (129 proteins in total) the score for position 21 is 13, which places it within, but towards the lower end of, the range of scores (9 to 24) for positions 5 to 30. The possibility that a positively-charged amino acid at either position 1, 2 or 3 in the N-terminal region of the signal peptide does not act as a signature component when accompanied by one or more positively-charged amino acids at position 4 or greater can be dismissed because, amongst the tester set of proteins, AvrSr35 and PGTG-08638 as well as the flax rust proteins AvrM-A, AvrL2-A and AvrL567-A, all have additional positively-charged amino acids at position 4 or greater (Fig. 1).

Finally there is a group of 106 proteins that do not possess a positively-charged amino acid at positions 1 or 2 or 3 but do possess one or more positively-charged amino acids at positions 4 or greater in the N-terminal region of the signal peptide. Here the score for position 21 (14) is undistinguished, sitting well within the range of scores (8 to 22) for the other positions (Table 1, row 9). This observation therefore provides no indication as to whether a positively-charged amino acid at position 4 or greater in the N-terminal region of the signal peptide might also act as a signature component.

In summary, the observations reported above clearly indicate that in wheat stem rust secreted proteins there is an association between the presence of a positively-charged amino acid at position 1 or 2 or 3 in the N-terminal region of the signal peptide and the frequency of positively-charged amino acids at position 21 downstream of the signal peptide relative to the frequency of positively-charged amino acids at other positions in the protein. This was the prediction if both components are required to mark a protein for translocation to the cytoplasm of the plant cell. Therefore this analysis supports the hypothesis that the translocation signature for rust secreted proteins consists of two components, one in the signal peptide and one downstream of the signal peptide. Given that two components, acting together, are required for translocation, it is suggested that an appropriate name for this signature would be the “Duet” signature.

The flax rust proteins in the tester group (Fig. 1), like the wheat rust proteins, all have a positively-charged amino acid at positions 2 and/or 3 in the N-terminal region of their signal peptides. However, unlike the wheat rust proteins, they do not possess a positively-charged amino acid at position 21 downstream of the signal peptide, but all do possess a positively-charged amino acid in the 17-20 range downstream of the signal peptide, this being the only four amino acid frame amongst the first 43 amino acids downstream of the signal peptide in which all seven flax rust proteins possess a positively-charged amino acid (Fig. 2). Assuming that there is a basic similarity between the translocation signatures of flax and wheat rust secreted proteins, it might be hypothesised that the flax rust translocation signature consists of a signal peptide component similar to that in the wheat rusts together with a positively-charged amino acid in the 17-20 range downstream of the signal peptide. The 17-20 range could be larger due to the small sample size (seven) on which it is based. It is assumed that the range in positions reflects the biological situation and has not arisen due to error variation made by the predictor in predicting the length of the signal peptide.

An analysis to examine whether there is an association between the two putative signature components in the flax rust secretome proteins, similar to that carried out for the wheat stem rust secretome proteins, was undertaken using a list of 851 proteins with signal peptides that were identified by the Joint Genome Institute following the sequencing of the flax rust genome [22]. This list was reduced to 524 proteins after the removal of those proteins with a predicted transmembrane domain (144), those whose amino acid sequence for the first 70 amino acids was identical to that of another protein adjacent to, or nearby, in the list (49) and those proteins for which PHOBIUS and/or SIGNALP 6.0 did not predict the presence of a signal peptide (134).

The final list of 524 proteins is much smaller than that used in the wheat stem rust analysis (1208), which reduces the effectiveness of the analysis. Only one (AvrL567-A) of the seven flax rust tester proteins (Fig. 1) occurs in the list of 524 proteins, which highlights its lack of comprehensiveness. By contrast, all three of the tester wheat stem rust proteins (Fig. 1) are included in the list of 1208 wheat stem rust proteins, although only the first 309 amino acids of the 578 amino acid AvrSr35 protein are listed.

With the flax rust secretome proteins it is the comparative frequency of positively-charged amino acids at positions 17-20 downstream of the signal peptide that is of interest. Similar to the wheat stem rust secretome proteins, amongst those proteins with no positively-charged amino acids in the N-terminal region of their signal peptides, the frequency of positively-charged amino acids at positions 17-20 is unexceptional, falling within the range set by the frequency at other positions (Table 2, row 1; Fig. 4A).

**Figure 4.**
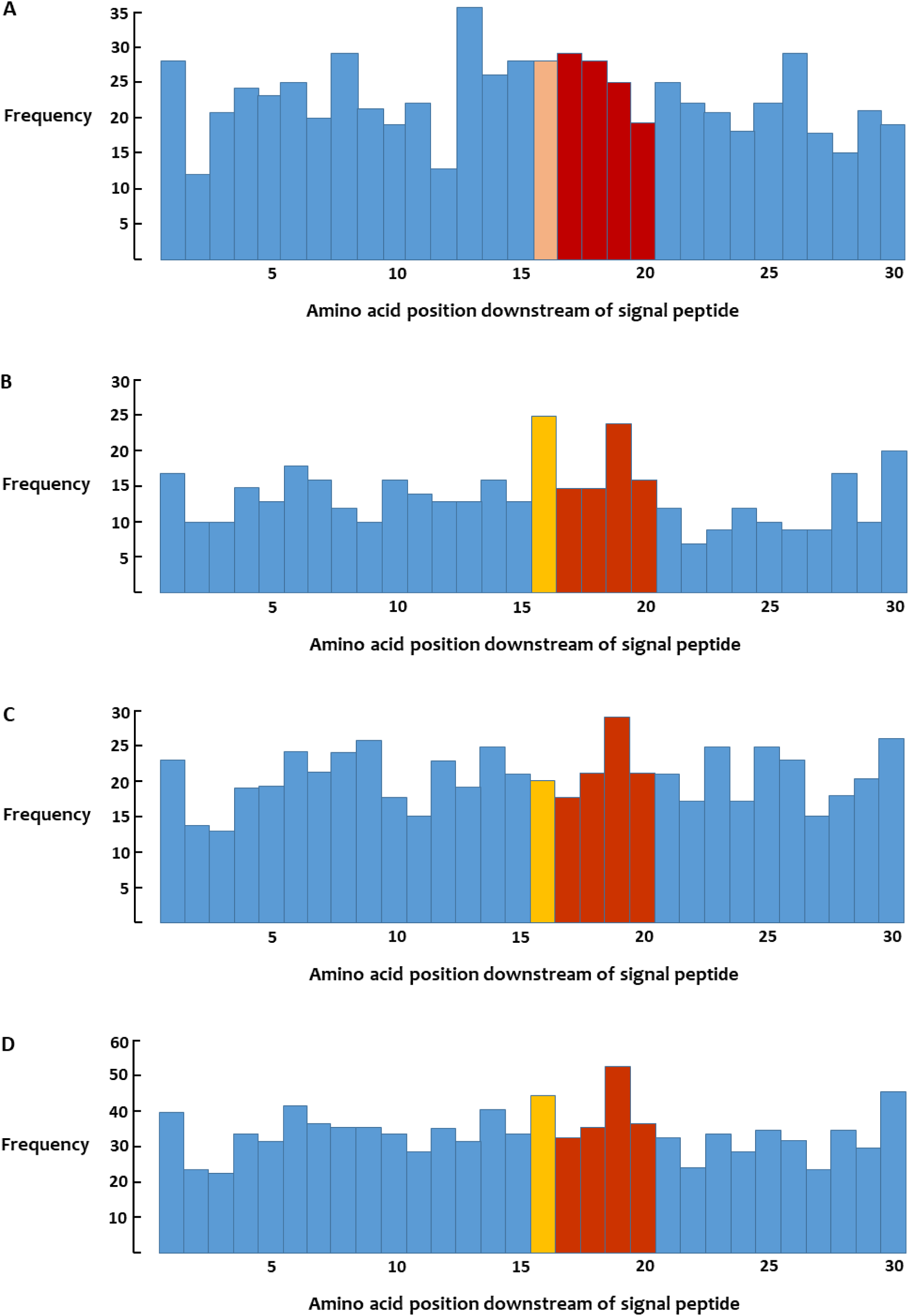
Frequency of positively-charged amino acids at positions 1 to 30 downstream of the signal peptide of flax rust secretome proteins. **A**. Group of 171 proteins that have no positively-charged amino acids upstream of the hydrophobic region in their signal peptides. **B**. Group of 107 proteins that have a single positively-charged amino acid at either position 2 or position 3 upstream of the hydrophobic region in their signal peptides with or without additional positively-charged amino acids at position 4 or greater. **C**. Group of 163 proteins that have a single positively-charged amino acid at position 1 upstream of the hydrophobic region in their signal peptides with or without additional positively-charged amino acids at position 4 or greater. **D**. Group of 270 proteins that have a single positively-charged amino acid at either position 1 or position 2 or position 3 upstream of the hydrophobic region in their signal peptides with or without additional positively-charged amino acids at position 4 or greater.

**Table 2.** Frequency of positively-charged amino acids at positions 1 to 30 downstream of the signal peptide of 524 flax rust secretome proteins grouped according to their positively-charged amino acid composition in the N-terminal region of their signal peptides

| Row | 1 | 2 | 3 | 4 | 5 | 6 | 7 | 8 | 9 | 10 | 11 | 12 | 13 | 14 | 15 | 16 | 17 | 18 | 19 | 20 | 21 | 22 | 23 | 24 | 25 | 26 | 27 | 28 | 29 | 30 |
| --- | --- | --- | --- | --- | --- | --- | --- | --- | --- | --- | --- | --- | --- | --- | --- | --- | --- | --- | --- | --- | --- | --- | --- | --- | --- | --- | --- | --- | --- | --- |
| 1 | 28 | 12 | 21 | 24 | 23 | 25 | 20 | 29 | 21 | 19 | 22 | 13 | 36 | 26 | 28 | 28 | 29 | 28 | 25 | 19 | 25 | 22 | 21 | 18 | 22 | 29 | 18 | 15 | 21 | 19 |
| 2 | 23 | 14 | 13 | 19 | 19 | 24 | 21 | 24 | 26 | 18 | 15 | 23 | 19 | 25 | 21 | 20 | 18 | 21 | 29 | 21 | 21 | 17 | 25 | 17 | 25 | 23 | 15 | 18 | 20 | 26 |
| 3 | 10 | 4 | 5 | 13 | 7 | 13 | 10 | 7 | 7 | 12 | 9 | 8 | 9 | 10 | 9 | 18 | 10 | 10 | 18 | 11 | 8 | 5 | 6 | 7 | 8 | 7 | 6 | 14 | 6 | 13 |
| 4 | 7 | 6 | 5 | 2 | 6 | 5 | 6 | 5 | 3 | 4 | 5 | 5 | 4 | 6 | 4 | 7 | 5 | 5 | 6 | 5 | 4 | 2 | 3 | 5 | 2 | 2 | 3 | 3 | 4 | 7 |
| 5 | 40 | 24 | 23 | 34 | 32 | 42 | 37 | 36 | 36 | 34 | 29 | 36 | 32 | 41 | 34 | 45 | 33 | 36 | 53 | 37 | 33 | 24 | 34 | 29 | 35 | 32 | 24 | 35 | 30 | 46 |
| 6 | 4 | 2 | 4 | 8 | 9 | 10 | 7 | 5 | 6 | 8 | 2 | 4 | 15 | 7 | 4 | 6 | 3 | 5 | 2 | 2 | 4 | 6 | 6 | 5 | 6 | 7 | 7 | 7 | 4 | 7 |
| 7 | 6 | 8 | 2 | 10 | 7 | 10 | 3 | 12 | 7 | 8 | 7 | 8 | 7 | 5 | 9 | 7 | 5 | 1 | 6 | 6 | 8 | 4 | 6 | 5 | 6 | 3 | 3 | 5 | 4 | 8 |

Row 1: No positively-charged amino acids in the N-terminal region of the signal peptide – group of 171 proteins
Row 2: Single positively-charged amino acid at position 1 upstream of hydrophobic region of signal peptide with or without additional positively-charged amino acids at position 4 or greater – group of 163 proteins
Row 3: Single positively-charged amino acid at position 2 upstream of hydrophobic region of signal peptide with or without additional positively-charged amino acids at position 4 or greater – group of 75 proteins
Row 4: Single positively-charged amino acid at position 3 upstream of hydrophobic region of signal peptide with or without additional positively-charged amino acids at position 4 or greater – group of 32 proteins
Row 5: Sum of rows 2, 3 and 4 – group of 270 proteins
Row 6: Two or three positively-charged amino acids at positions 1, 2 and 3 upstream of hydrophobic region of signal peptide with or without additional positively-charged amino acids at position 4 or greater – group of 40 proteins
Row 7: No positively-charged amino acid at position 1, 2 or 3 upstream of hydrophobic region of signal peptide but have one or more positively-charged amino acids at position 4 or greater – group of 43 proteins

Amongst those proteins with a positively-charged amino acid at position 2 in the signal peptide N-terminal region, with or without additional positively-charged amino acids at position 4 or greater (75 proteins), 18 possess a positively-charged amino acid at position 19, which is equal highest with position 16, followed by position 28 with 14 (Table 2, row 3). For those proteins with a positively-charged amino acid at position 3 in the signal peptide N-terminal region, with or without additional positively-charged amino acids at position 4 or greater (32 proteins), 6 possess a positively-charged amino acid at position 19, which is exceeded by two positions, one of which is position 16 (Table 2, row 4). When these two groups are combined, positions 16 and 19 stand out, having frequencies of 25 and 24 respectively, with the next highest scores being 20 and 18 for positions 30 and 6 respectively (Fig. 4B).

Amongst those proteins with a positively-charged amino acid at position 1 in the signal peptide N-terminal region, with or without additional positively-charged amino acids at position 4 or greater (a larger sample of 163 proteins), position 19 again stands out with a score of 29, while the next highest score amongst positions 5-30 is 26 at position 9 (Table 2, row 2; Fig. 4C).

Thus, as with the wheat stem rust secretome proteins, the signal peptide position 1 group is distinguished by the same characteristic as the position 2 and position 3 groups, suggesting again that a positively-charged amino acid at position 1 in the signal peptide should also be included in the hypothesised component 1 of the translocation signature.

When the three groups above are combined (270 proteins), the score for position 19 is 53, with position 30 in second place with a score of 46, followed by position 16 with a score of 45 (Table 2, row 5; Fig. 4D). Thus position 19 consistently stands out. Furthermore, the sum of the scores for positions 17-20 when the three groups are combined (Table 2, row 5; Fig. 5A) is 159, which is exceeded only by the sum of scores for positions 16-19 (167) because position 16 consistently scored high values, possibly indicating that position 16 is also a signature position. In support of this, for AvrM14-A, PHOBIUS, SIGNALP 3.0, 4.1 and 5.0, all predict a signal peptide of 20 amino acids but SIGNALP 6.0, as used in the main analysis, predicts a 21 amino acid signal peptide, thereby placing a positively-charged amino acid at position 16, instead of position 17, downstream of the signal peptide. The sum of the scores for positions 16-20 (three groups combined – Table 2, row 5; Fig. 5B) is 204, which is greater than the sum of the scores for any other five amino acid consecutive set not involving positions 16-20, which ranged from 154-185 for positions 5-30. Thus this analysis of flax rust secretome proteins, although limited by the size of the data set, does point to an association between the presence of a positively-charged amino acid at positions 1, 2 or 3 upstream of the hydrophobic region in the signal peptide and the comparative frequency of a positively-charged amino acid at positions (16)17-20, particularly position 19, downstream of the signal peptide.

**Figure 5.**
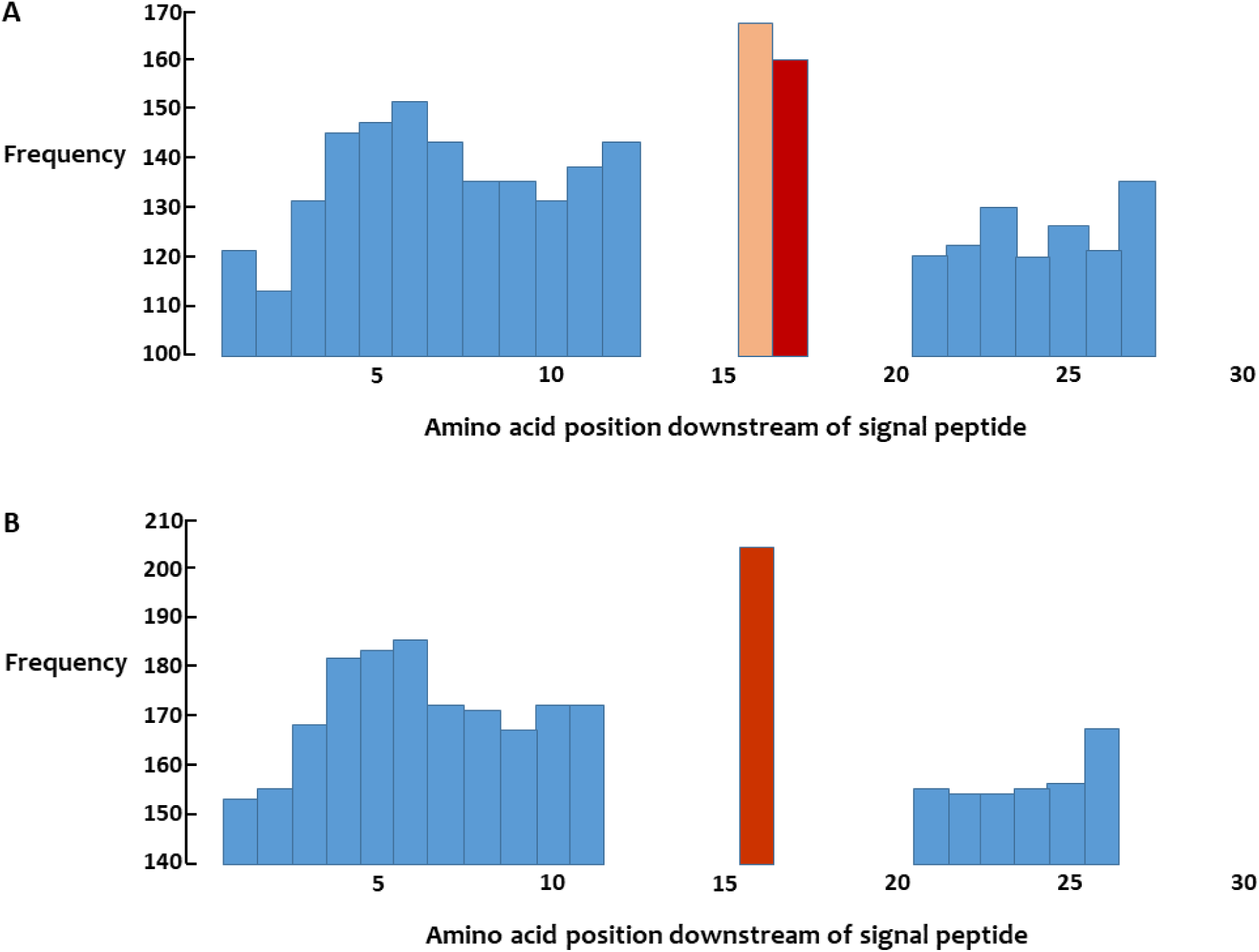
Frequency of positively-charged amino acids at four or five consecutive positions at positions 1 to 30 downstream of the signal peptide of 270 flax rust secretome proteins that have a positively-charged amino acid at position 1 or 2 or 3 upstream of the hydrophobic region in the signal peptide with or without additional positively-charged amino acids at position 4 or greater. **A**. Frequency of four consecutive positions beginning at position shown. **B**. Frequency of five consecutive positions beginning at position shown.

Thus the flax rust data analysis also supports a two-component translocation signature but with the second component (a positively-charged amino acid at positions 17-20, possibly 16-20) differing from that indicated in the wheat stem rust analysis (a positively-charged amino acid at position 21). Interestingly, like in the flax rust proteins, position 19 also stands out in the wheat stem rust proteins where it also had the highest score for the number of positively-charged amino acids amongst 723 secretome proteins that have one or more positively-charged amino acids at position 1 or 2 or 3 in the N-terminal region of the signal peptide with or without additional positively-charged amino acids at position 4 or greater (position 19 score = 124 cf. a range of 86-110 for positions 5-30, excluding position 21 – see Table 1, row 8 and Fig. 6). This suggests the possibility that wheat stem rust might be employing two translocation signatures. One, like flax rust, where the second component consists of a positively-charged amino acid at position (16) 17-20 downstream of the signal peptide; this could be employed during the rust’s pycnial stage when it is growing on its alternate dicot host, barberry.

**Figure 6.**
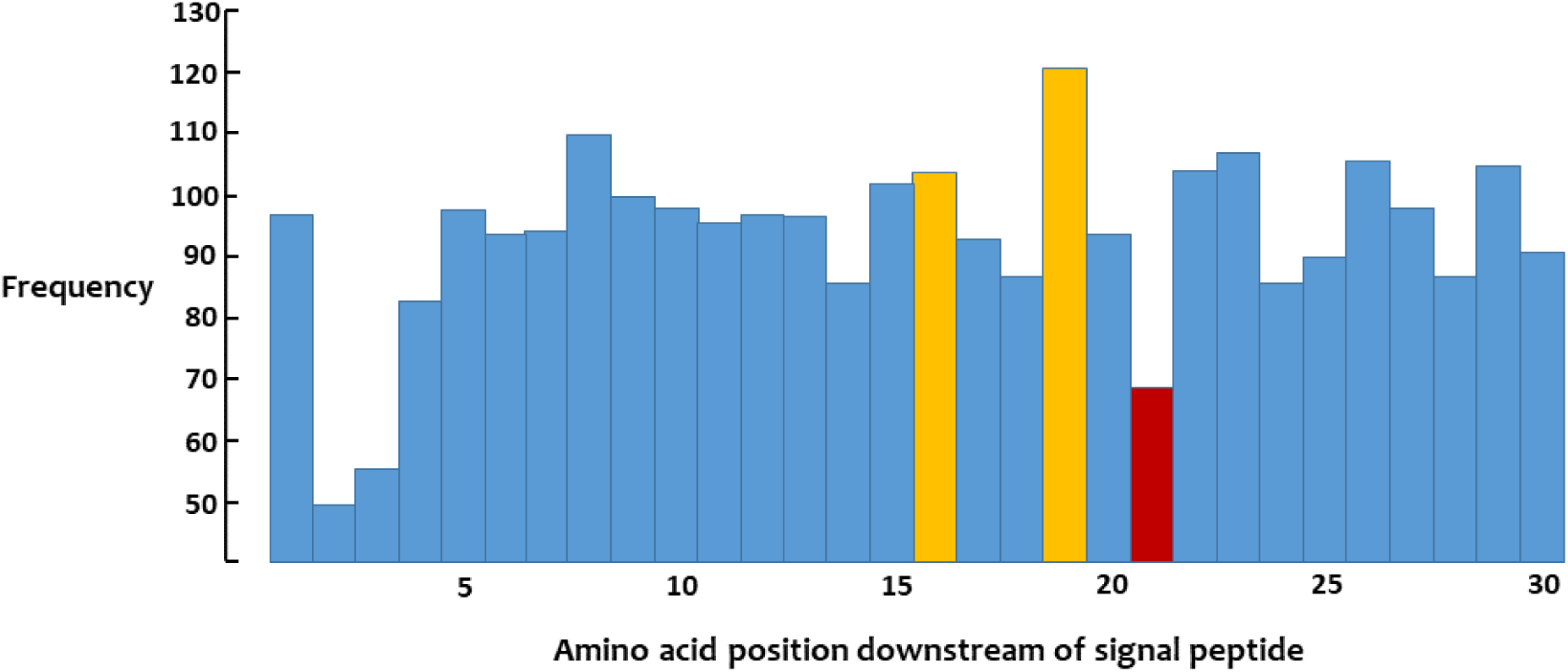
Frequency of positively-charged amino acids at positions 1 to 30 downstream of the signal peptide amongst a group of 723 wheat stem rust secretome proteins that have one or more positively-charged amino acids at position 1 or 2 or 3 upstream of the hydrophobic region in their signal peptides with or without additional positively-charged amino acids at position 4 or greater.

The second translocation signature, involving a positively-charged amino acid at position 21 (the +21 duet signature), could then subsequently have evolved for use during the rust’s asexual stage when growing on its monocot host, wheat. Many of the effectors that facilitate growth on the dicot host are likely to be different from those that facilitate growth on the monocot host and having different translocation signatures would be a way of achieving the necessary specificity.

In the oomycete pathogens, where an RxLR motif in the N-terminal region of the protein marks a protein for translocation to the cytoplasm of the plant cell, there is also a (previously un-noted) association with the presence of a positively-charged amino acid in the N-terminal region of the signal peptide. Raffaele et al. [23, see additional file 1] identified 2228 *Phytophthora infestans* proteins that contain a secretion signal peptide (using four different predictors) whose amino acid sequences were obtained from the National Centre for Biotechnology information (general portal, *Phytophthora infestans*, strain T30.4, genome assembly ASM14294v1). The first 1327 proteins on this list were examined for the presence of an RxLR motif and submitted to the PHOBIUS predictor. The list of 1327 proteins was reduced to 763 proteins after removal of (i) those proteins with a predicted transmembrane domain (ii) those proteins that were repeats (for the first 90 amino acids) of other proteins and (iii) those proteins for which PHOBIUS and/or at least one of the other four predictors did not predict a signal peptide. Amongst the 763 proteins, 206 contain an RxLR motif (R+), and 557 do not (R-). In the signal peptide, PHOBIUS found that 612 contain one or more positively-charged amino acids in the N-terminal region of their signal peptides (SP+) while 151 do not (SP-). The joint segregation data are R+SP+ (191), R+SP- (15), R-SP+ (421) and R-SP- (136). Thus 191 of the 206 proteins with an RxLR motif also possess a signal peptide with one or more positively-charged amino acids in their N-terminal region. The 2x2 contingency table χ^2^ value for the data is 26.75 (P < 0.001). Thus there is a highly-significant association between the presence of an RxLR motif and the presence of a positively-charged amino acid in the N-terminal region of the signal peptide which, in 159 of the 191 instances, included a positively-charged amino acid, usually arginine, adjacent to the initiating methionine. If the 15 proteins with the RxLR motif that do not have a positively-charged amino acid in the N-terminal region of their signal peptide do not get translocated to the cytoplasm of the plant cell then it could be hypothesised that *P. infestans* also has a two-component translocation signature.

## Discussion

There was not a high level of agreement between the predictions of PHOBIUS and those of SIGNALP 6.0. Their predictions for the size of the signal peptide differed on 326 occasions amongst the 1208 wheat stem rust proteins analysed, a difference that affects the positioning of positively-charged amino acids downstream of the signal peptide. However, a larger and more significant disagreement occurred in the prediction of the size of the N-terminal region within the signal peptide, where the predictors differed on 808 occasions, with SIGNALP 6.0 predicting a larger size on 113 occasions and a smaller size on 695 occasions. In this latter group the larger N-terminal region PHOBIUS prediction contained one or more positively-charged amino acids in 555 instances: consequently, using the smaller SIGNALP 6.0 predictions would have resulted in a change in the number and/or positioning of the positively-charged amino acids in the N-terminal region in each of these 555 proteins which would have greatly affected the analysis. If the SIGNALP 6.0 predictions had been used in the tester set of proteins (Fig. 1), the signal peptide N-terminal region signature component would not have been identified because in nine of the ten proteins listed in Fig. 1 (SIGNALP 6.0 did not predict a signal peptide for AvrP) the SIGNALP 6.0 prediction for the size of the N-terminal region in the signal peptide was smaller than that predicted by PHOBIUS, thereby completely changing the distribution of the positively-charged amino acids in this region and, in the case of Pst_8713, predicting an N-terminal region with no positively-charged amino acids. Therefore, in any future analysis of rust secretome proteins the PHOBIUS predictor should definitely be used to predict the size of the N-terminal region in the signal peptide. Given that SIGNALP 6.0 predictions (long output, slow model mode) for the size of the signal peptide leads to all four of the wheat rust tester proteins having a positively-charged amino acid at position 21 (as in Fig. 2 where SIGNALP 4.1 was used) and resulted in an association being detected between the signal peptide N-terminal region signature component and the comparative frequency of a positively-charged amino acid at position 21, it would seem sensible to use the size predictions of this predictor.

Recently three related genes encoding secreted proteins at the *AvrSr27* locus in wheat stem rust strain Pgt21-0 were identified [24]. Two, *AvrSr27-1* and *AvrSr27-2*, were located on the chromosome associated with avirulence on host lines possessing *Sr27* while the other, *avrSr27-3*, was located on the chromosome associated with virulence on *Sr27*, as determined by deletion mutations. However, when recombinant *Barley stripe mosaic virus* (BSMV), expressing each of the *AvrSr27* genes without their predicted signal peptides, were tested on host lines possessing *Sr27* it was found that all three were unable to infect, whereas they could infect host lines without *Sr27*. The observation that *avrSr27-3,* while having a virulent phenotype in wheat stem rust, has an avirulent phenotype in the BSMV test, was ascribed as likely to be due to expression levels as *avrSr27-3* was found to account for only 15% of the total expression of the *AvrSr27* locus in isolated haustoria, with expression of *AvrSr27-2* being four fold higher than both *AvrSr27-1* and *avrSr27-3*. It was suggested that, in the absence of *AvrSr27-1* and *AvrSr27-2*, the expression of *avrSr27-3* was below a threshold required to confer avirulence on host lines with *Sr27*. It was also suggested that *AvrSr27-2* likely contributes the most to avirulence during infection of *Sr27* plants. With regard to the duet signature, all three of the proteins have a positively-charged amino acid immediately following the initiating methione which, in AvrSr27-2 and avrSr27-3, are located at position 4 upstream of the hydrophobic region in the signal peptide based on SIGNALP 6.0 predicting an N-terminal region of five amino acids. PHOBIUS does not predict a signal peptide for any of the proteins so its predictions, which have been used in all previous analyses, cannot be use here. Concerning the second component of the duet signature, an examination of the amino acid sequence of the three proteins shows that, given a signal peptide of 28 amino acids as determined by SIGNALP 5.0 (and by SIGNALP 6.0 for AvrSr27-2 and avrSr27-3), only AvrSr27-2 has a positively-charged amino acid at position 21 downstream of the signal peptide. Therefore it is predicted that only AvrSr27-2 is translocated into the plant cell where it is recognised by the intracellular plant resistance protein Sr27, a coiled coil-nucleotide binding site-leucine rich repeat protein [24]. On this basis, AvrSr27-2 is solely responsible for the avirulence phenotype of wheat stem rust strain Pgt21-0 on *Sr27* plants.

In the current analysis 69 wheat stem rust secretome proteins were found to possess both components of the proposed translocation signature (a positively-charged amino acid at position 1 and/or 2 and/or 3 in the N-terminal region of the signal peptide with or without additional positively-charged amino acids at position 4 or greater, together with a positively-charged amino acid at position 21 downstream of the signal peptide – see Table 1, rows 5, 6 and 7). This should facilitate the identification of avirulence proteins since those recognised by intracellular resistance proteins should all be in this group. This group of 69 could be larger as 269 proteins were excluded from the analysis because PHOBIUS and/or SIGNALP 6.0 did not predict the presence of a signal peptide (even though SIGNALP 4.0 did) and many of these are likely to be genuine secreted proteins. Here it might be noted that the signal peptides of two of the proteins in the tester set (Fig. 1), AvrP123 and AvrP4, were not predicted by PHOBIUS and SIGNALP 6.0 respectively. Also, there is still uncertainty as to whether a positively-charged amino acid at position 4 or greater upstream of the hydrophobic region in the signal peptide acts as a signature component. Thus 69 should be seen as the lower limit of the number of proteins translocated to the cytoplasm of the plant cell. Proteins translocated to the cytoplasm of the plant cell are highly likely to “alter plant processes” and, therefore, to be effectors. Of the 69 “signature” proteins, 55 contain less than 400 amino acids and only three contain more than 650 amino acids.

In the flax rust analysis of 524 proteins, 130 possess both components of the translocation signature assuming a 17-20 frame for the position of the positively-charged amino acid downstream of the signal peptide or 156 proteins for a 16-20 frame. If the sequencing of the flax rust genome had been as comprehensive as that of the wheat stem rust genome and an equivalent number of secretome proteins had been identified (1208), these figures of 130 and 156 would proportionally increase to 300 and 360 respectively. These numbers are considerably larger than the 69 found to possess both components amongst 1208 wheat stem rust proteins. A possible reason for this difference is that, with flax rust, the sexual and asexual cycles occur on the same host plant (autoecious). So flax rust most probably has had a continuous association, potentially for up to 175 million years when the early angiosperms arose [25], with a single host, even though that host will have evolved over time, thereby providing a long time period for cytoplasmic effectors to evolve. By contrast, with wheat stem rust, the sexual and asexual cycles occur on different host species (heteroecious), with the jump to a monocot for the asexual stage likely occurring sometime after the grass family (*Poaceae*) proliferated at the end of the Cretaceous period (145-66 Mya) or during the Miocene (23-5.3 Mya) when grasses gained importance in southern Europe and Asia Minor [26]. Thus wheat stem rust likely has had a lot less time to evolve cytoplasmic effectors adapted to its asexual cycle host compared to flax rust. Also, as noted above, wheat stem rust could have a separate set of cytoplasmic effectors adapted to its dicot host (barberry) that have a flax rust-like translocation signature. If so, 334 of the 1208 wheat stem rust secretome proteins have both translocation signature components assuming a 17-20 range for the second component and 389 for a second component range of 16-20, figures similar to those predicted for the flax rust genome. If the flax rust-like signature (the +(16-20) duet signature) does operate in wheat stem rust when growing on barberry then, amongst the 69 proteins predicted to be translocated when the rust is growing on wheat, 37 are also predicted to be translocated when growing on barberry, since these have positively-charged amino acids in the 16-20 range as well as at position 21. If some of the genes encoding secretome proteins are expressed only in the sexual stage, and others only in the asexual stage, then the number of proteins being transported into a plant cell would be less than the figures above.

Concerning the mechanism of translocation, there are two broad possibilities. It could be a two-step process, whereby the protein is first secreted into the extrahaustorial matrix where it is then “recognised” and incorporated into a mechanism that moves it into the plant cell. Alternatively, it could be a one step process, whereby those proteins with a translocation signature are recognised soon after their synthesis, perhaps even before they have folded, and incorporated into a separate secretion and translocation pathway, such as secretion into a specialised vesicle. The second possibility would seem the more likely, given that one component of the translocation signature is located in the signal peptide and, under the two step model, a signal peptide with special characteristics would not be required. Oomycetes likely possess a separate secretion and translocation pathway, given the finding that secretion and translocation of an RxLR containing cytoplasmic effector from *P. infestans* is not inhibited by brefeldin A, which inhibits conventional Golgi-mediated secretion [27].

A study of the RxLR containing protein Avr3a from the oomycete pathogen, *P. infestans*, has shown that the RxLR motif is cleaved off prior to secretion by the pathogen with the N-terminus of the mature effector likely to be acetylated [28]. The possibility that a similar cleavage might occur in rust proteins targeted to the host cytoplasm at the signature positively-charged amino acid at position 21 (wheat stem rust) or positions (16)17-20 (flax rust) is not supported by a study of the structure of the flax rust avirulence protein AvrP [29]. This study found that the AvrP protein binds three Zn ions and that each Zn ion has a tetrahedral coordination with either four cysteine residues or three cysteine and one histidine residue. Relevant here is the finding that two of the cysteine residues coordinating one Zn ion occur at positions 13 and 15 downstream of the signal peptide, just upstream of the two signature positively-charged amino acids at positions 16 and 17. It would seem unlikely that these two critical cysteine residues would be cleaved off.

Recently, in work aimed at identifying additional avirulence genes in wheat stem rust, a library of wheat stem rust genes encoding secreted proteins was screened by transforming them into plant protoplasts containing various wheat stem rust resistance genes. Library-specific RNA-seq and differential gene expression analysis that identified reduced expression levels was taken as an initial indication that the pathogen gene had triggered a resistance response. Additional functional tests were then carried out to confirm that a resistance response occurs when the putative avirulence gene and the corresponding resistance gene are present in the same cell. In this way two wheat stem rust genes, *AvrSr13* and *AvrSr22*, were identified that interact with the wheat rust resistance genes *Sr13* and *Sr22,* respectively [30].

An examination of the amino acid sequences of the protein products of *AvrSr13* and *AvrSr22* reveals that neither of them possess the hypothesised duet signature, an observation that calls into question the validity of the duet signature hypothesis. Relevant here is the observation that all the avirulence genes whose protein products do possess the hypothesised duet signature have two-factor authentication in that a candidate gene was first identified, either because it co-segregated with a known avirulence gene in a mapping family (as with the flax rust avirulence genes), or because it was associated with a loss-of-avirulence mutation event (as with the wheat stem rust avirulence genes *AvrSr27*, *AvrSr35* and *AvrSr50*). Subsequently functional tests confirmed that the candidate gene triggered a resistance response when present in the same cell as the corresponding plant resistance gene. In contrast, *AvrSr13* and *AvrSr22* only have single-factor authentication in that they were directly identified by a functional test. This functional test assumes that the resistance gene protein only interacts with a single protein in the rust secretome. If, however, the resistance gene protein has the capability to interact with more than one protein in the rust secretome (as has recently been shown with Sr27 [31]), then another (apoplastic) protein might be incorrectly identified as the avirulence protein. Consequently, the lack of the duet signature in the AvrSr13 and AvrSr22 proteins should not, at this stage, be cause for rejecting the duet signature hypothesis.

It should be noted that the duet signature identified in wheat stem rust that operates during the asexual cycle (the +21 duet signature) is not necessarily expected to be found in other *Puccinia* species growing on monocot hosts since these other *Puccinia* species that have different alternate hosts would have followed separate evolutionary pathways in developing the ability to grow on monocot hosts. This applies to wheat leaf rust (*Puccinia triticina*), whose alternate host is *Thalictrum speciosissimum* or *Isopyrum fumaroides*, and southern corn rust, *Puccinia polysora*, whose alternate host is unknown. The one wheat leaf rust avirulence gene, *AvrLr15*, [32] and two southern corn rust avirulence genes, *AvrRppC* and *AvrRppK*, [33,34] identified to date all lack the +21 duet signature – all three genes were identified by direct functional tests so come with the qualification noted above. Wheat stripe rust may employ the +21 duet signature since it has the same alternate host as wheat stem rust and one of its proteins with the +21 duet signature was included in the tester set of proteins (Fig. 1). If the +21 duet signature is operative in wheat stripe rust then it can be hypothesised that the +21 duet signature initially arose in a barberry rust species and that this species subsequently diverged into wheat stem rust and wheat stripe rust. It will be of interest to determine whether *P. triticina*, *P. polysora* and other monocot rusts have also evolved separate translocation signatures specific to their monocot hosts or whether they have adapted the dicot +(16-20) duet signature for use when growing on monocot hosts.

Rusts are highly sophisticated parasites as might be expected after an association of some 200 million years with their plant hosts [21]. At the molecular level very little is known about their “modus operandi” in three key areas. First, to what extent, if at all, does the rust influence the lipid and protein composition of the extrahaustorial membrane. Second, what is the function of each of the large numbers of proteins secreted by haustoria, not only those that are translocated to the cytoplasm of the plant cell but also those that are secreted into the extrahaustorial matrix. Third, what is the mechanism that translocates proteins from the cytoplasm of the rust cell into the cytoplasm of the plant cell. This mechanism is likely to be unique to rusts and its elucidation would be a major advance in our understanding of rust biology and could, potentially, reveal new approaches to controlling rust disease. The identification of the rust protein translocation signature should assist work in this area and also, by identifying which secreted rust proteins go to the plant cell cytoplasm and which to the extrahaustorial matrix, provide guidance in determining the function of a secreted protein.

## Acknowledgements

I thank Narayana Upadhyaya for directions to the list of flax rust secreted proteins and Evans Lagudah for comments on an earlier version of this manuscript.

